# The microbial tryptophan metabolite indole acts on the gastrointestinal tract to improve glucose homeostasis by enhancing GLP-1 secretion and L-cell differentiation

**DOI:** 10.1101/2025.06.24.661421

**Authors:** Phyllis Phuah, Mariana Norton, Sijing Cheng, Anna G. Roberts, Leah Meyer, Pei-En Chung, Cecilia Dunsterville, Rafal Karwowski, Brian Y.H. Lam, Emile Otsubo, Sofia Aleksashina, Fiona M. Gribble, Frank Reimann, Aylin C. Hanyaloglu, Giles S.H. Yeo, Gavin A. Bewick, Ben Jones, Bryn Owen, Kevin G. Murphy

**Affiliations:** Section of Endocrinology and Investigative Medicine, Division of Diabetes, Endocrinology and Metabolism, Department of Metabolism, Digestion, and Reproduction, Faculty of Medicine, 6th Floor Commonwealth Building, Hammersmith Hospital, London, W12 0NN, UK; MRC Metabolic Diseases Unit, Institute of Metabolic Science Metabolic Research Laboratories, University of Cambridge, Cambridge, CB2 0QQ, UK; Institute of Reproductive and Developmental Biology, Department of Metabolism, Digestion and Reproduction, Imperial College London, London, W12 0NN, UK; Diabetes and Obesity Theme, School of Cardiovascular and Metabolic Medicine and Sciences, Faculty of Life Sciences and Medicine, King’s College London, SE1 1UL, UK

**Keywords:** glucagon-like peptide-1, glucose homeostasis, transient receptor potential channel subtype A1

## Abstract

**Aims/hypothesis:** Growing evidence implicates gut microbiota-derived metabolites in metabolic homeostasis. Indole, a microbial tryptophan metabolite, has been reported to enhance Glucagon-like peptide-1 (GLP-1) secretion in vitro, and its derivatives have been inversely associated with risk of type 2 diabetes (T2D). We hypothesised that indole acts via the gastrointestinal tract to modulate glucose homeostasis, and aimed to test this hypothesis using in vitro and in vivo models.

**Methods:** The acute effects of indole on GLP-1 secretion *in vitro*, and on glucose tolerance and hormone secretion in mice, were determined. Subsequently, the effects of indole on intestinal epithelial cell fate and L-cell differentiation in murine ileal organoids and *in vivo* were studied. Finally, the utility of chronic indole administration in a murine model of T2D was explored.

**Results:** Indole stimulated *in vitro* GLP-1 secretion in a concentration-dependent manner, and improved acute glucose control *in vivo.* Additionally, we demonstrate that indole drives enteroendocrine L-cell differentiation in murine ileal organoids, resulting in increased L-cell density and longer-term glucoregulatory benefits *in vivo.* Finally, sub-chronic indole administration improved glucose tolerance and insulin sensitivity in diabetic mice.

**Conclusions/interpretation:** Our findings identify indole as an anti-diabetic molecule that acts on the gut, and raise the possibility of incorporating indole into nutraceutical supplements to aid in the treatment or prevention of T2D. This highlights the importance of gut microbiota-derived metabolites in metabolic health and opens new avenues for developing novel strategies to combat T2D.

**Research in Context:** *What is already known about this subject?:*

- Gut microbiota-derived metabolites play a role in metabolic homeostasis.
- Indole, a microbial tryptophan metabolite, enhances GLP-1 secretion *in vitro*.
- Indole derivatives are inversely associated with type 2 diabetes (T2D) risk.

*What is the key question?:*

- Does indole act on the gut to modulate glucose homeostasis?

*What are the new findings?:*

- Indole stimulates GLP-1 secretion and improves acute glucose control *in vivo*.
- Indole drives enteroendocrine L-cell differentiation in murine ileal organoids, increasing L-cell density and long-term glucoregulatory benefits.
- Sub-chronic indole administration improves glucose tolerance and insulin sensitivity in mice with type 2 diabetes, showing the potential of gut microbiota-derived metabolites as therapeutic targets.

*How might this impact clinical practice in the foreseeable future?:*

- Indole could be incorporated into nutraceutical supplements for T2D prevention or treatment.

## 1. Introduction

The gastrointestinal tract plays a crucial role in maintaining metabolic and glucose homeostasis. Beyond its role in digestion and absorption, it regulates systemic metabolism and glucose levels through modulation of gut hormone secretion and the gut microbiome^1,2^. This intricate relationship between the gut, its microbial inhabitants, and host metabolism is essential for health and has implications for metabolic disorders.

Gut microbial dysbiosis is implicated in the pathophysiology of metabolic disorders like obesity and type 2 diabetes (T2D). Nutrient-derived metabolites are key mediators of host-microbiota crosstalk and hence metabolic homeostasis^3^. An inverse association between plasma concentrations of derivatives of indole, a bacterial catabolite of the essential amino acid L-tryptophan, and T2D risk was recently identified^4^. While most dietary tryptophan is absorbed for protein synthesis or used as a substrate for kynurenine and serotonin pathways, 4-6% is metabolised by colonic bacteria to generate indole and its derivatives. Indole is the most abundantly generated microbial tryptophan catabolite, with concentrations of 0.2-1mM detected in human and rodent faeces^5^. Locally, indole enhances intestinal epithelial integrity and attenuates inflammation^6,7^. However, recent studies have also reported that indole modulates gut hormone release. Specifically, indole was found to stimulate glucagon-like peptide-1 (GLP-1) secretion from GLUTag cells following short exposures in vitro^8,9^. GLP-1 is an incretin secreted by enteroendocrine L-cells. Synthetic GLP-1 analogues are current T2D pharmacotherapies, though their utility can be limited by side-effects. Accordingly, an alternative treatment strategy is to modulate endogenous GLP-1 release with dietary metabolites. We report for the first time that indole improves acute glucose tolerance in vivo and drives enteroendocrine L-cell differentiation, and that sub-chronic indole administration improves glucose tolerance and insulin sensitivity in both healthy and diabetic mice.

## 2. Materials and Methods

### 2.1 Animals

Mice were eight to twelve weeks old and specific pathogen free. Mice weighed 20-32g, apart from TALLYHO mice, which weighed 30-40g. Male C57Bl/6J mice (Charles River, UK) were used, unless otherwise stated. Mice on a 129svJ background were bred in-house. GluCreERT2 mice were kindly gifted by Fiona Gribble and Frank Reimann and crossed to Ai95(RCL-GCaMP6f)-D mice (JAX strain: #028865). GLP-1R fl/fl mice were kindly gifted by Professor Randy Seeley at University of Michigan and crossed with Pdx1-Cre/Esr1 mice (JAX strain: #024968). Tamoxifen (100mg/kg) was injected intraperitoneally for 4-days to induce beta cell knockout of the GLP-1R. Both male and female GluCreERT2;Ai95-D and Pdx1-Cre;GLP-1R fl/fl mice were used. Only male TALLYHO/Jng mice, purchased from the Jackson Laboratory (stock: #005314, JAX, USA), were used because only the males develop diabetes. A total of 388 mice were used.

All mice were group housed in individually ventilated cages containing standard bedding on a 12h-light/dark cycle (lights on 07:00, lights off 19:00) with ad libitum access to standard laboratory RM1 chow and water and cardboard tubes for enrichment. Procedures were carried out in the light cycle. All procedures were carried out according to the Animals (Scientific Procedure) Act, 1986 and approved by Imperial College London Animal Welfare and Ethical Review Body. Our study followed ARRIVE reporting guidelines. All manipulations used mice from different groups (randomised by body weight) in a mixed sequence to avoid confounders, and investigators were blinded.

### 2.2 Intraperitoneal Glucose Tolerance Test

Glucose tolerance tests were performed on mice after a 5h fast with ad libitum access to water. Following baseline blood glucose measurement, mice were intraperitoneally injected with 20% glucose (2g/kg). Changes in blood glucose were measured over 120 minutes from tail-tip venesection using an AccuCheck glucometer. Glucose was administered intraperitoneally rather than orally to bypass the gastrointestinal tract, allowing us to study glucose clearance and insulin secretion independent of glucose-stimulated insulin release and any potentially confounding effects on gastric emptying.

To investigate the effect of indole on acute glucose tolerance, indole (20mg/kg; I3408, Sigma-Aldrich, UK) or vehicle (water) was orally gavaged 15 minutes before glucose IP. This dose of indole was selected based on earlier studies showing an effect of indole on reducing mucosal inflammation^10^.

Fasting blood glucose levels varied significantly in the TALLYHO mice due to the different extent of diabetes progression in each mouse. The raw IPGTT data was therefore normalised to t0 baseline for the TALLYHO studies to determine whether indole could improve glucose homeostasis in this model.

### 2.3 Food Intake Study

Tests were performed in the early light phase on individually caged mice following a 16h fast. Mice received an oral gavage of indole (20mg/kg) or vehicle (water) following which pre-weighed amounts of RM1 chow was returned to the cages. Changes in food intake were measured over 2h.

### 2.4 Intraperitoneal Insulin Tolerance Test

Insulin tolerance tests were performed on mice after a 5h fast with ad libitum access to water. Following baseline blood glucose measurement, mice were injected with insulin at 1 IU/kg (Actrapid, Novo Nordisk, Denmark). Changes in blood glucose were measured over 120 minutes from tail-tip venesection using an AccuCheck glucometer.

### 2.5 Static glucose-stimulated insulin secretion assay

Pancreatic islets were isolated from adult C57Bl6/J mice by collagenase digestion as previously described^11^ and maintained at 37°C, 5% CO_2_ in RPMI medium with 10% FBS and 1% penicillin-streptomycin overnight.

Islets were incubated at 37°C for 1 hour in HEPES Krebs Ringer bicarbonate buffer (HKRB) (pH 7.4, bubbled with CO2) with 1% BSA and 3mM glucose (i.e., low glucose). Next, the islets were stimulated with HKRB with 11mM glucose (i.e., high glucose) in the presence or absence of 1mM indole. Supernatant samples (which contain secreted insulin) and lysed islet samples were separately collected and stored at −20°C until further use. Insulin levels were assayed using the mouse Insulin ELISA (Cisbio Bioassays, Codolet, France) according to the manufacturer’s instructions.

### 2.6 Gut hormone secretion assays

All cell and organoid lines were routinely tested for mycoplasma contamination. STC-1 cells were maintained in DMEM with 10% FBS, 1% pen-strep at 37°C, 5% CO2. Colonic crypts were isolated from 6-8 week old C57Bl6/J mice as previously described^12,13^ and plated onto 24-well, 2% Matrigel-coated plates. The crypts were cultured in DMEM (with 10% FBS, 1% pen-strep and 10μM Y-27632) and incubated overnight at 37°C, 5% CO_2_.

Hormone secretion assays were performed using STC-1 cells and primary murine colonic crypt cultures. The cells were treated with test reagents (made up in DMEM) for 2h at 37°C, 5% CO_2_. Supernatant and lysed cell samples were collected and stored at −20°C. GLP-1 and PYY levels in cell supernatants and lysates, and in plasma, were measured using previously described sensitive and specific in-house radioimmunoassays^14,15^.

### 2.7 Ileal organoid cultures

Ileal organoids were derived from the last 6 cm of the ileum of 6-12 week old C57Bl6/J and PPG-Cre;GCaMP6 mice, as previously described^16^. Dissociated crypts were resuspended in Cultrex Pathclear Reduced Growth Factor Basement Membrane Extract (BME) and cultured in Intesticult media at 37°C with 5% CO2, with medium changes every 2-3 days. Organoids were passaged every 5-7 days in a 1:2-1:3 ratio using Gentle Cell Dissociation Reagent and mechanical disruption.

### 2.8 Organoid imaging experiments

PPG-Cre;GCaMP6 organoids were seeded onto 96-well black-walled plates. After three days, they were treated with 1μM of 4-hydroxytamoxifen for 24 hours to induce GCaMP6f expression in L-cells. Timelapse recordings were used to assess calcium response following addition of treatment. Forskolin (10μM) and ATP (100μM), were added after 10 minutes to confirm cell viability. Imaging was performed using a Nikon ti2e widefield microscope, capturing 1 frame per second.

Images were analyzed using ImageJ with a custom macro package (Intensity_2) from Imperial FILM facility. Regions of interest (ROIs) were drawn around cell bodies to measure fluorescence intensity (f). Baseline fluorescence was calculated from 10-30 seconds prior to treatment addition. Fluorescence change was quantified as Δf/fbaseline, with the maximum value calculated over 10 minutes post-treatment. Cells were then stratified based on the following cut off – responder: max Δ f/fbaseline of treatment greater than 20% of Fmax-Fmin^17^.

### 2.9 Organoid immunostaining

Murine ileal organoids for imaging were plated onto 8-well chamber slides (Thermofisher Scientific, UK). After treatment, organoids were fixed with 4% PFA, permeabilised with 0.1% Triton-X PBS, and blocked with 5% BSA for 1h at room temperature. Primary antibody incubation was performed overnight at 4oC using mouse anti-GLP-1 antibody (ab23468, 1:500; Abcam, UK). Subsequently, the organoids were washed and incubated with the secondary antibody anti-mouse Alexa Fluor 647-conjugated IgG (1:1000; ThermoFisher, UK) for 1h at RT. DAPI was used as a nuclear stain. Images were acquired using a Leica SP5 confocal laser scanning microscope.

### 2.10 Tissue processing and immunostaining

IHC was performed per standard protocols. Tissues were fixed overnight in 4% PFA at 4°C and processed into paraffin blocks. Following sectioning using a Leitz 1512 microtome, 5μm sections were deparaffinised, rehydrated, and subjected to antigen retrieval via heating to 95°C in 10 mM sodium citrate buffer for 40 mins, followed by incubation in 2M HCl for 20 min, and 0.05% trypsin solution for 15 min at 37°C. Next, sections were blocked in 5% BSA (Sigma-Aldrich, UK) and 5% normal goat serum (Sigma-Aldrich, UK) for 2h at RT. Primary antibody incubation was performed overnight at 4°C using mouse anti-GLP-1 antibody (ab23468, 1:500; Abcam, UK). Sections were washed and incubated with the secondary antibody anti-mouse Alexa Fluor 647-conjugated IgG (1:1000; ThermoFisher, UK) for 1h at RT. DAPI was used to counterstain the nuclei. Tile scans of the ileum sections were obtained using a Nikon Eclipse Ti2E-2 widefield microscope. The number of GLP-1 immunoreactive cells in stained organoid and ileum samples was quantified manually using the Cell Counter plugin in ImageJ and calculated as a proportion of DAPI-stained nuclei. The counter was blinded to treatment.

### 2.11 RNA isolation for gene expression analysis by qPCR and sequencing

Organoids were cultured and treated as described above and snap frozen at −80°C. Total RNA was isolated using TRIsure (phenol)-chloroform extraction after homogenisation with TissueLyser II. Following cDNA synthesis using a High Capacity cDNA Reverse Transcription Kit (Applied Biosystems) according to the manufacturer’s instructions, qPCR was performed using Maxima Probe Master Mix (Life Technologies, Paisley, UK) and Taqman PCR core reagents (Applied Biosystems, UK) in a C1000 CFX384 Real-Time System Thermocycler. Data were analyzed using the 2^−ΔΔCt^ method.

For bulk RNA sequencing, the right liver lobe was dissected from mice and snap frozen at −80°C. Total RNA was purified by using the RNeasy® Plus Mini kit (Qiagen, Hildon, Germany) according to the manufacturer’s instructions.

### 2.12 RNA sequencing

Purified total RNA samples were analysed using nanodrop (A260/280 >1.8) and the RNA integrated were checked using Agilent Bioanalyzer RNA 6000 Nano assay (RIN >7). Strand-specific mRNA (polyA) libraries were constructed at Novogene UK Ltd and sequenced using Illumina NovaSeq 600 with 5.8-7.9 billion bases per sample. Adapters were trimmed using Novogene’s bioinformatics pipeline and alignment were performed using HISAT2^18^ and Mouse genome GRCm38 Ensembl version 102. Uniquely mapped reads were taken forward for gene-level counts and differential gene expression analysis using DESeq2^19^.

### 2.13 Pathway analysis

The dataset was submitted for Ingenuity Pathway Analysis (http://www.ingenuity.com/). A core analysis was conducted using a cutoff p value of <0.05 which reduced the analysis ready dataset to 716. Stringent filters (Benjamini-Hochberg p < 0.05, z-score > 2) were applied to detect enriched canonical pathways. For upstream regulator analysis, a z-score cutoff of >2 and a p value of overlap <0.05 was used.

### 2.14 Statistical analysis

Data were analysed using Prism 9.0 (GraphPad) and are expressed as mean ± SEM. Student’s t-test was used for two-group comparisons. One-way ANOVA with Tukey’s post-hoc was used for single independent variable experiments with >2 groups. Two-way ANOVA with Tukey’s post hoc was used for two independent variables. Significance was set at p < 0.05.

## 3. Results

### 3.1 Indole improves acute glucose tolerance by potentiating GLP-1 secretion via Trpa1

An oral indole pre-load significantly improved acute intraperitoneal glucose tolerance (Figure 1A) but an intraperitoneal indole pre-load did not (Fig 2A), suggesting the effects of indole are mediated via the gut, potentially by modulating gut hormones. Furthermore, other tryptophan metabolites did not have similar effects on glucose tolerance. Administration of the major tryptophan metabolite kynurenic acid via oral gavage or intraperitoneal injection, at doses previously shown to affect inflammasome activation in colitis^22^, did not affect glucose tolerance (Fig 4A, B). Although previous studies have reported chronic indole suppresses food intake^20,21^, we did not identify any acute effects of indole on food intake or insulin sensitivity in lean mice (Fig 2B, C).

**Figure 1.**
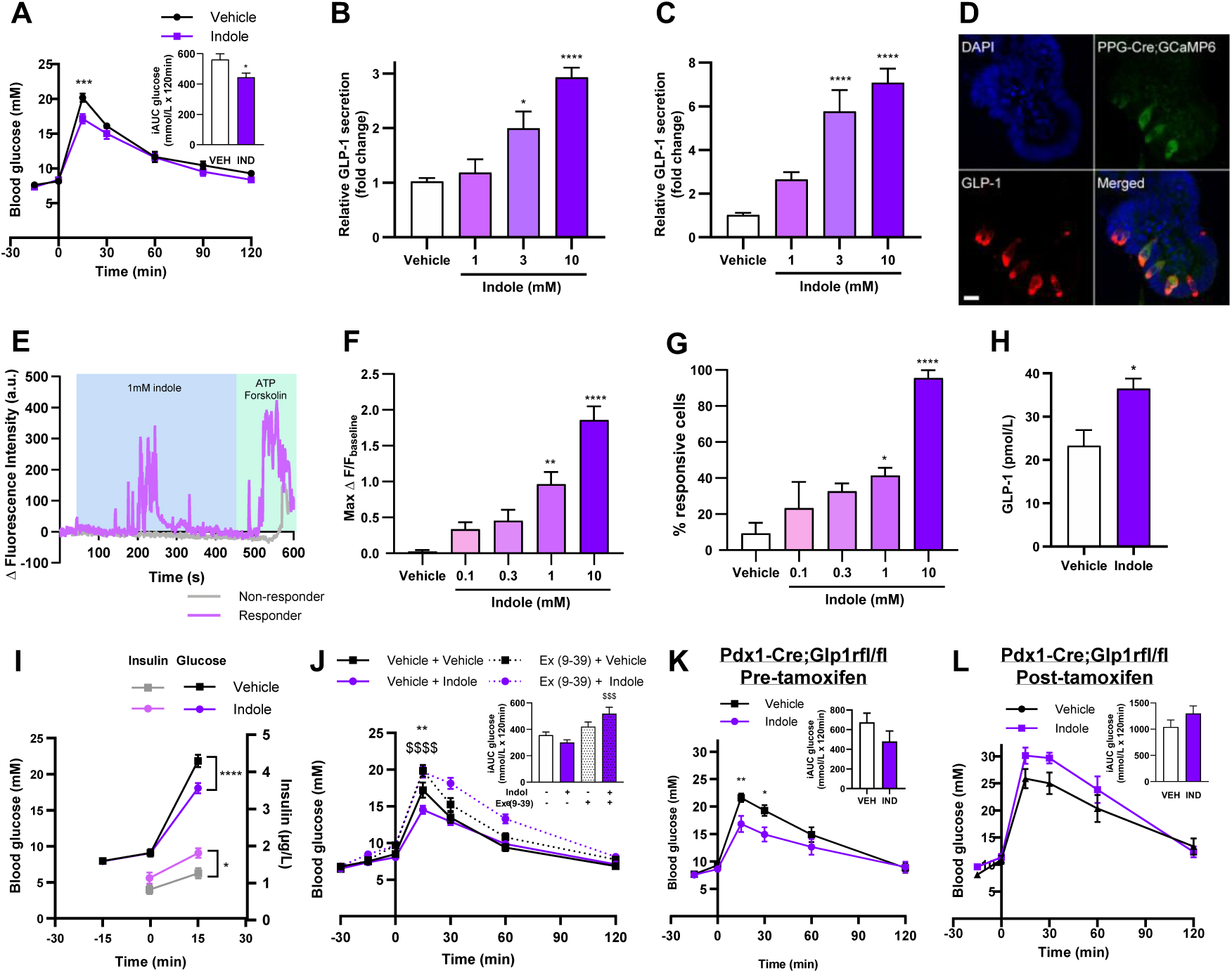
Indole potentiates GLP-1 secretion to improve acute glucose tolerance. (A) Blood glucose excursion during an intraperitoneal glucose tolerance test (IPGTT) after a 5h fast in lean C57BL6/J mice orally gavaged with indole (20mg/kg) or water vehicle 15 minutes prior to glucose load (2g/kg). n=8-10/group. The effect of indole (1-10mM) on GLP-1 secretion from (B) STC-1 cells and (C) primary murine colonic crypts, following a 2h incubation. n=3-8 independent plates. (D) GLP-1 immunostaining of 4-OHT treated PPG-Cre;GCaMP6 murine ileal organoids to induce the expression of GCaMP6 in L-cells. (E) Representative temporal traces of [Ca^2+^] changes in individual L-cells following 1mM indole treatment of PPG-Cre;GCaMP6 murine ileal organoids, expressed as Δ Fluorescence intensity. ATP and forskolin were applied at the end of each imaging session as a positive control. (F) The effect of indole (0.1-10mM) on the magnitude of L-cell calcium response. (G) The proportion of L-cells showing a calcium response following indole (0.1-10mM) treatment. n=19-90 cells from 7-33 organoids. (H) The effect of an oral gavage of indole (20mg/kg) or water vehicle on plasma GLP-1 concentration 15 minutes after administration in C57Bl6/J mice following a 5h fast. n=6/group. (I) C57Bl/6 mice were pre-treated with an oral gavage of indole (20mg/kg) or water vehicle 15 minutes prior to intraperitoneal glucose injection (2g/kg). Blood glucose levels (vehicle—black, indole—purple) and plasma insulin levels (vehicle—grey; indole—light purple) were measured at t-15 (for glucose), t0 and t15 timepoints. n=8-9/group. (J) Blood glucose excursion during a IPGTT in C57BL6/J mice that received an intraperitoneal injection of the GLP-1 receptor antagonist exendin (9-39) or saline vehicle 30 minutes, and an oral gavage of indole (20mg/kg) or water vehicle 15 minutes prior to glucose load. n=12-14/group. Pdx1-CreERT;Glp1rfl/fl mice were generated following tamoxifen administration (100mg/kg for 5 consecutive days) to induce knockdown of GLP-1R in Pdx1-expressing β-cells. Blood glucose excursion during IPGTT (K) before and (L) after tamoxifen treatment. Indole (20mg/kg) or water vehicle was administered via oral gavage 15 min prior to intraperitoneal glucose bolus (2g/kg). n=5-6/group. Insets in (A), (J), (K), (L), (H) represent incremental area under the curve (iAUC) relative to t0. (A), (I)-(L), analysed by two-way ANOVA with Tukey’s post-hoc. (B), (C), (F)-(H) analysed by one-way ANOVA with Tukey’s post-hoc. (H) and iAUCs in (A), (J), (K), (L) analysed by student’s t-test. Error bars represent SEM. *p<0.05, **p<0.01, ***p<0.001, ****p<0.0001, $$$$p<0.0001 vehicle+indole vs Ex(9-39)+indole.

**Figure 2.**
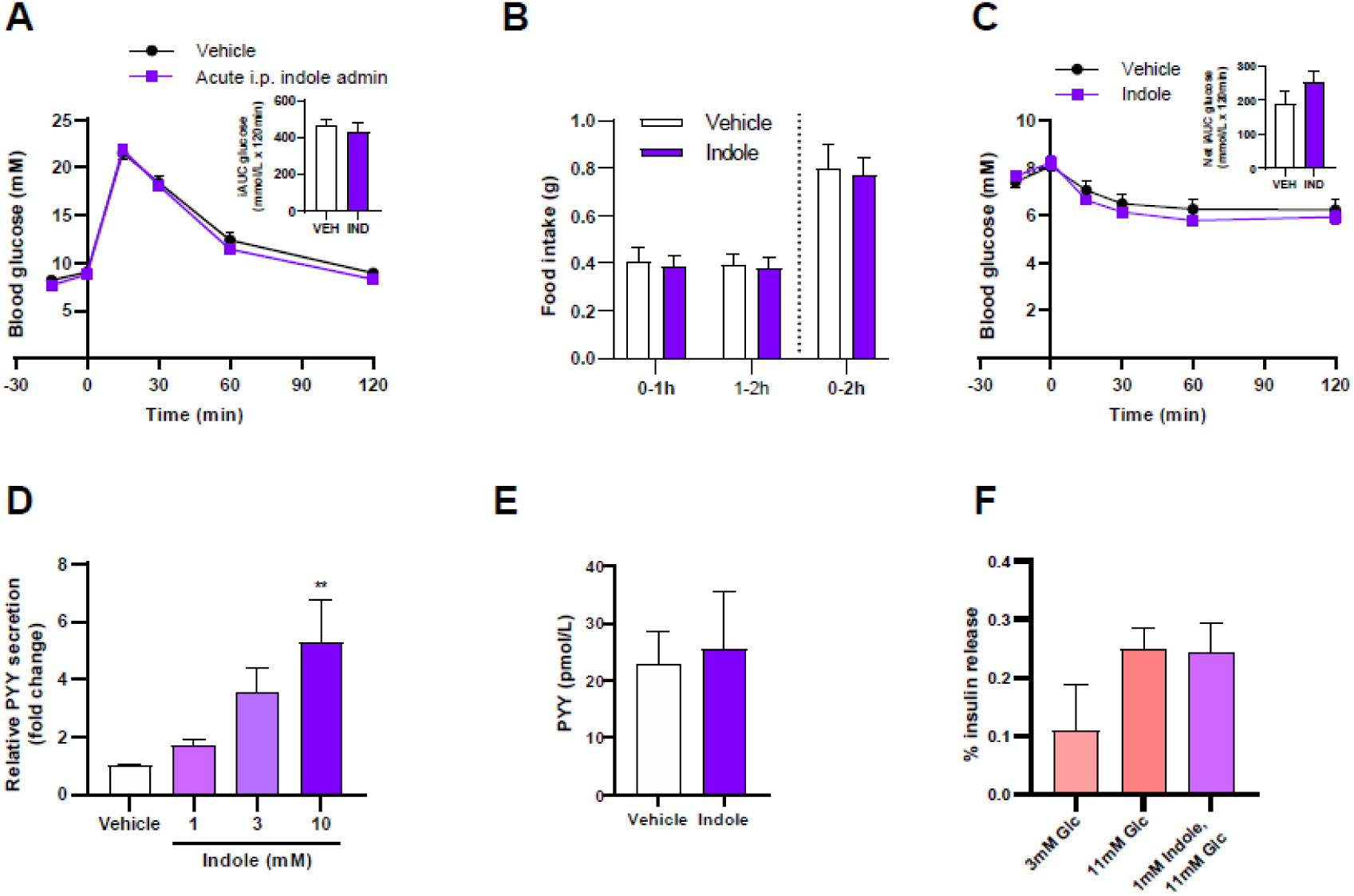
Indole does not modify acute food intake, and does not improve acute glucose tolerance via direct pancreatic action. (A) Blood glucose excursion during IPGTT in C57BL6/J mice that received an intraperitoneal injection of either indole (20mg/kg) or water vehicle 15 minutes prior to glucose load. n=10/group. Inset represents iAUC determined relative to t0. (B) The effect of indole (20mg/kg) oral gavage or water vehicle on food intake in overnight fasted male mice at the onset of the early light phase for 0-2H post-administration. n=9-10/group. (C) Blood glucose excursion during an intraperitoneal insulin tolerance test after a 5h fast in lean C57BL6/J mice that received an oral gavage of 20mg/kg indole or water vehicle 15 minutes prior to insulin injection (1IU/kg). n=8-9/group Inset represents iAUC determined relative to t0. (D) The effect of indole (1-10mM) on PYY secretion from primary murine colonic crypts, following a 2h incubation. n=3-8 independent plates. (E) The effect of an oral gavage of indole (20mg/kg) or water vehicle on plasma PYY concentrations 15 minutes after administration in C57Bl6/J mice following a 5h fast. (F) The effect of 1mM indole in the presence of high glucose (11mM) on insulin secretion from dispersed murine pancreatic islets, following 1h static incubation. N=3 independent experiments. (A), (B), (C) analysed by two-way ANOVA with Tukey’s post-hoc. (D), (F) analysed by one-way ANOVA with Tukey’s post-hoc. (D) and iAUCs in (D), (E) analysed by student’s t-test. Error bars represent SEM. **p<0.01.

Given previous reports that indole modulates gut hormone secretion in vitro, we hypothesised that the acute glucoregulatory effects of indole were mediated via enhanced GLP-1 secretion. We found that 10mM indole increased GLP-1 release ∼3-fold from the enteroendocrine STC-1 cell line (Fig 1B), while both 3mM and 10mM indole increased GLP-1 release ∼6-fold from murine colonic L-cell cultures (Fig 1C). Live cell imaging on transgenic organoids generated from PPG-Cre;GCaMP6 murine ileal crypts, in which L-cells express the fluorescent Ca2+ indicator GCaMP6f (Fig 1D), showed that indole induced a rapid and reversible dose-dependent elevation of intracellular Ca2+, with 10mM indole increasing maximum fluorescence intensity two-fold and the percentage of responsive L-cells (Fig 1E-G). Correspondingly, oral administration of indole in mice significantly increased plasma total GLP-1 levels after 15 minutes (Fig 1H). Together, this extends previous in vitro findings^8,9^ that indole stimulates GLP-1 release. Furthermore, following oral indole administration in mice, insulin secretion increased in parallel with the improvement in glucose tolerance (Fig 1I). However, indole did not potentiate insulin secretion from murine islets in vitro (Fig 2F), suggesting it does not directly modulate pancreatic hormone secretion. The acute glucose-lowering effects of oral indole administration were dependent on GLP-1 signalling. These effects were significantly attenuated by pre-treatment with the GLP-1 receptor antagonist exendin (9-39) (Fig 1J). There was no longer a beneficial effect of indole on glucose tolerance in Pdx1-CreERT;Glp1r fl/fl transgenic mice, in which GLP-1 receptor expression is inducibly knocked down in Pdx1-expressing pancreatic beta cells (Fig 1K, 1L). Together, these data suggest that the acute glucoregulatory effects of indole are mediated by increased GLP-1 release driving insulin secretion.

Although 10mM indole increased release of peptide YY (PYY), co-expressed with GLP-1 in L cells, by ∼4-fold from murine colonic crypt cultures (Fig 2D), plasma PYY levels were unchanged 15 minutes following oral indole administration in mice (Fig 2E). This is in accord with the lack of effect of indole on food intake in vivo, and suggests any effect on PYY release in vivo was likely short-lived. It may also reflect the relatively low density of PYY-expressing cells in the proximal small intestine.

Indole and its derivatives can modulate intestinal homeostasis and immune functions via the aryl hydrocarbon receptor (AhR)^5^, while impaired AhR ligand production has been associated with an increased risk of developing metabolic syndrome^24^. Given that many AhR antagonists have partial agonist actions, and can exhibit different efficacy depending on the specific ligand being blocked and the tissue the receptor is expressed in, we investigated whether AhR agonism might play a role in glucoregulation. However, at concentrations and doses previously shown to stimulate hormone secretion and reduce obesity-induced hyperglycaemia^24^, the synthetic AhR agonist Ficz did not stimulate GLP-1 secretion from STC-1 cells (Fig 4D), nor did Ficz modify glucose tolerance in C57Bl6/J mice (Fig 4C). This suggested that the AhR was not involved in the acute effects of indole on glucoregulation and GLP-1 secretion, perhaps unsurprisingly given that indole is a relatively weak agonist of the mouse AhR^25^. The excitatory calcium-permeable cation channel transient receptor potential channel subtype A1 (Trpa1) is widely expressed in EECs, and has been implicated in gut microbial EEC activation in zebrafish, as well as in indole-mediated serotonin secretion from human and murine primary intestinal cultures^9^. Trpa1 activation has also has previously been linked to GLP-1 secretion from murine L-cells *in vitro*^26^. Indeed, we found indole-induced GLP-1 release from STC-1 cells (Fig 3A) and murine primary colonic crypt cultures (Fig 3B) was attenuated in the presence of the Trpa1 inhibitor HC030031, while pre-treatment of PPG-Cre;GCaMP6 organoids with HC030031 blocked indole-stimulated L-cell activation (Fig 3C, D). When PPG-Cre;GCaMP6 organoids were imaged in Ca^2+^ free buffer, indole-stimulated Ca^2+^ mobilisation was blocked, reflecting the necessity of extracellular Ca^2+^ influx for indole to activate L-cells (Fig 3F, G), in accord with the action of Trpa1. Additionally, Trpa1 inhibition in mice also blocked indole-mediated improvements in glucose tolerance (Fig 3E).

**Figure 3.**
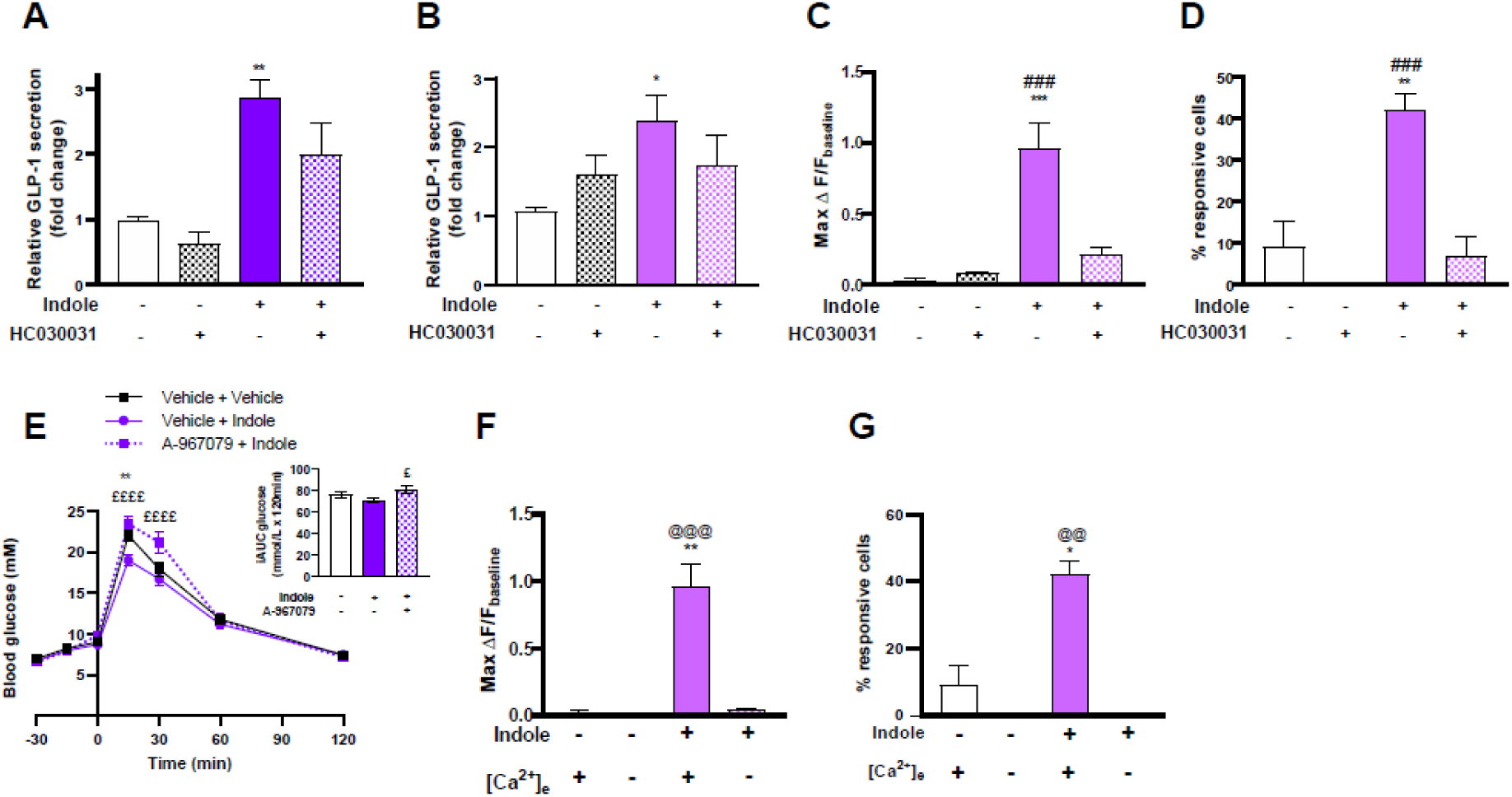
Indole modulates GLP-1 secretion via Trpa1 activity. The effect of indole in the presence or absence of the Trpa1 antagonist HC030031 on GLP-1 release from (A) STC-1 cells and (B) primary murine colonic crypts following a 2h incubation. n=4-7 independent plates. (C) The effects of 1mM indole in the presence or absence of HC030031 on L-cell calcium response of PPG-Cre;GCaMP6 organoids. (D) The proportion of L-cells showing calcium mobilisation following 1mM indole treatment in the presence or absence of 100μM HC030031. n=11-90 cells from 5-33 organoids. (E) Blood glucose excursion during a IPGTT in C57BL6/J mice that received an intraperitoneal injection of the Trpa1 inhibitor A-967079 (100mg/kg) or 6% DMSO, 4% Tween-20 in saline vehicle 30 minutes, and an oral gavage of indole (20mg/kg) or water vehicle 15 minutes prior to glucose load. n=10-12/group. Inset represents iAUC relative to t0. (F) The effects of 1mM indole in the presence or absence of extracellular Ca2+ in imaging buffer on L-cell calcium response of PPG-Cre;GCaMP6 organoids. (G) The proportion of L-cells showing calcium mobilisation following 1mM indole treatment in the presence or absence of extracellular Ca2+ in imaging buffer. n=16-78 cells from 6-30 organoids. (A)-(D), (F), (G) analysed by one-way ANOVA with Tukey’s post-hoc. (E) analysed by two-way ANOVA with Tukey’s post-hoc. iAUC in (E) analysed by student’s t-test. Error bars represent SEM. *p<0.05, **p<0.01, ***p<0.001, ###p<0.001 vehicle+indole vs HC030031+indole, $$$$p<0.0001 vehicle+indole vs Ex(9-39)+indole, ££££p<0.0001 vehicle+indole vs A-967079+indole. @@p<0.01 indole+ [Ca^2+^]_e_ vs. indole+no [Ca^2+^]_e_, @@@p<0.001 indole+ [Ca2+]e vs. indole+no [Ca^2+^]_e_.

The indole precursor molecule L-tryptophan can also stimulate GLP-1 release, an effect thought to be mediated in part via the Ca^2+^ sensing receptor^27^. While L-tryptophan induced Ca^2+^ mobilisation in L-cells was expectedly attenuated by the Ca^2+^ sensing receptor antagonist NPS2143, it was unmodified by HC030031 (Fig 4E, F). Together, these data suggest the Trpa1 ion channel mediates indole-stimulated GLP-1 secretion and glucose control via a process distinct from that which mediates tryptophan-induced GLP-1 secretion.

**Figure 4.**
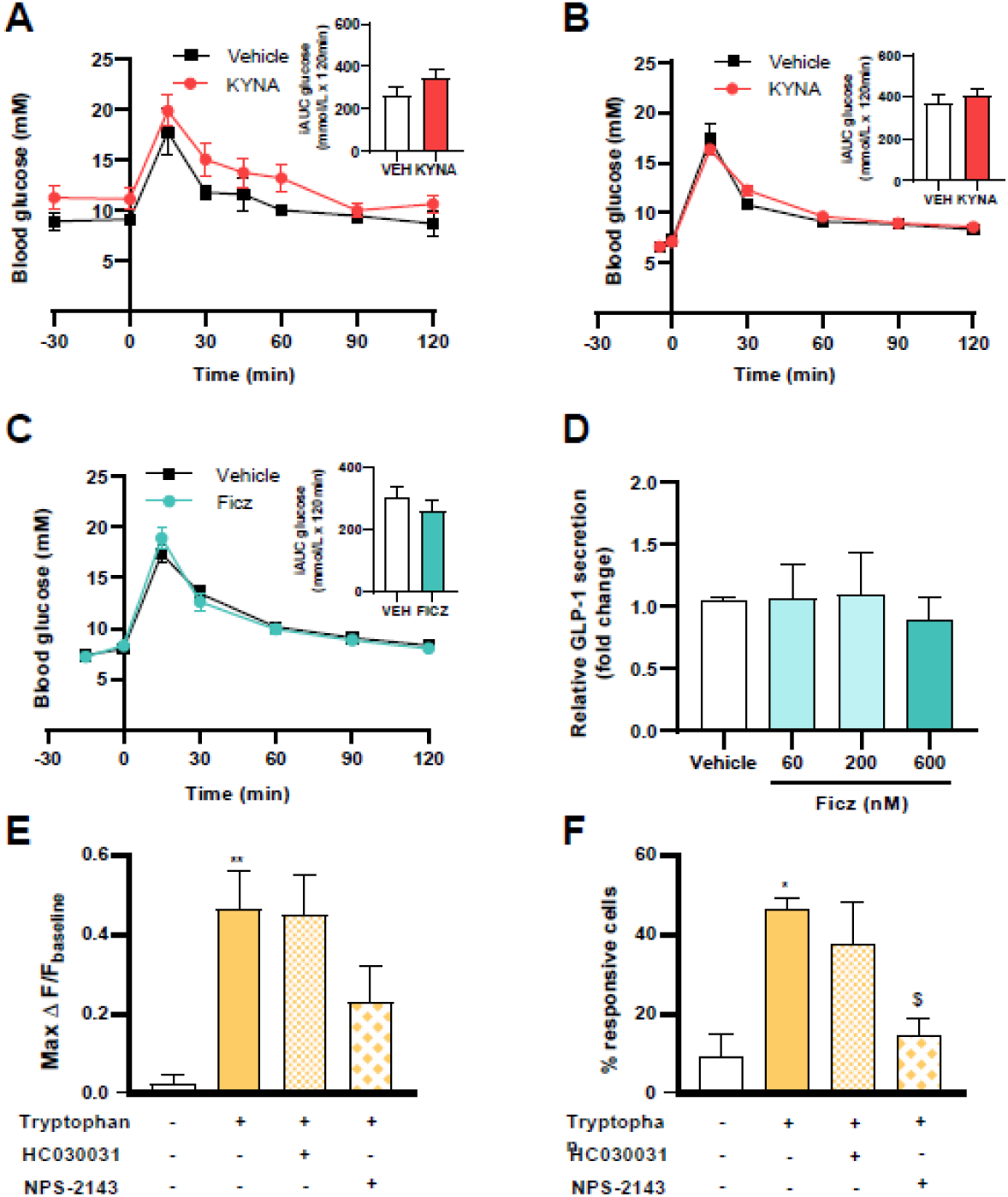
Investigating the effects of indole-related molecules on glucose homeostasis. (A) Blood glucose excursion during IPGTT in C57BL6/J mice that received an oral gavage of either 10mg/kg kynurenic acid or water vehicle 30 minutes prior to glucose load. n=7/group. Inset represents iAUC determined relative to t0. (B) Blood glucose excursion during IPGTT in C57BL6/J mice that received an intraperitoneal injection of either 5mg/kg kynurenic acid or water vehicle 5 minutes prior to glucose load. n=8/group. Inset represents iAUC determined relative to t0. (C) Blood glucose excursion during IPGTT in C57BL6/J mice that received an oral gavage of either Ficz (1µg) or 1% DMSO vehicle 15 minutes prior to glucose load. n=8-10/group. Inset represents iAUC determined relative to t0. (D) The effect of Ficz (60, 200, 600nM) on GLP-1 secretion from STC-1 cells, following a 2h incubation. n=6 independent plates. (E) The effects of 30mM tryptophan in the presence or absence of 100μM HC030031 or NPS2143 on L-cell calcium response of PPG-Cre;GCaMP6 organoids. (F) The proportion of L-cells which show calcium mobilisation following 30mM tryptophan treatment in the presence or absence of 100μM HC030031 or NPS2143. n=25-39 cells from 8-13 organoids. (A)-(C) GTTs analysed by two-way ANOVA with Tukey’s post-hoc, iAUCs analysed by student’s t-test. (D)-(F) analysed by one-way ANOVA with Tukey’s post-hoc. Error bars represent SEMs. *p<0.05, **p<0.01, $p<0.05 tryptophan+vehicle vs tryptophan+NPS2143.

### 3.2 A single bolus of indole improves long-term glucose tolerance by augmenting enteroendocrine differentiation and hence L-cell number

Specific GLP-1 secretagogues can enhance L-cell differentiation^28^. Furthermore, the tryptophan derivative indole-3-acetate has also been shown to modify intestinal lineage differentiation^29^. Having established the acute effects of indole on glucoregulation and GLP-1 secretion, we investigated whether indole mediated longer term effects on the GLP-1 system. Wildtype murine ileal organoids were exposed to a 3h treatment pulse of different concentrations of indole and the effect on expression of Gcg, which encodes GLP-1, determined 2 days later (Fig 6E). We subsequently repeated this study using the most effective concentration of 100µM indole (Fig 5A). We found a 1.24-fold upregulation in mRNA expression of Gcg in indole-treated organoids (Fig 5B), in addition to elevated transcript levels of Pyy and Gip (Fig 6A). Indole treatment also increased mRNA expression of the transcription factors NeuroD1 and Ngn3, which are associated with L-cell endocrine specification (Fig 5B), indicating enhanced development of EE secretory progenitors. The enteroendocrine differentiation factors Pax4 and Pax6 expression were also non-significantly raised (Fig 6A). In line with the upregulated Gcg mRNA expression, addition of 100µM indole to organoid culture medium for 3h increased L-cell number 1.55-fold (Fig 5C-E). Thus a short indole treatment exposure can augment the L-cell population in mouse intestinal organoids.

**Figure 5.**
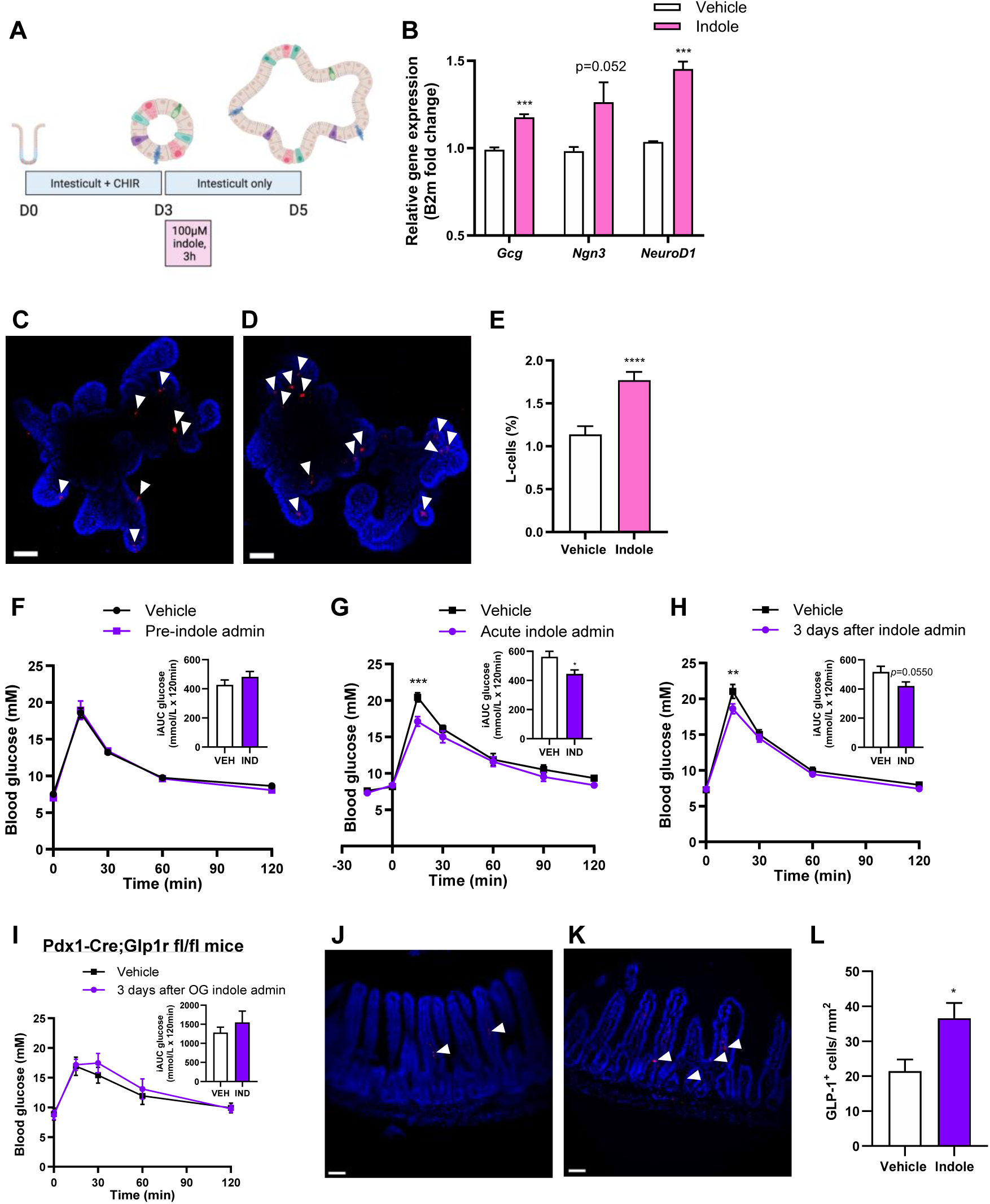
Indole increases L-cell abundance to improve long-term glucose tolerance. (A) Experimental paradigm for treatment of WT murine ileal organoids. Organoids were incubated with 100μM of indole or vehicle control for 3h and collected 2 days later. (B) Gene expression of the gut hormone Gcg, and the transcription factors directing L-cell development, Ngn3, NeuroD1,in murine intestinal organoids cultured with indole or vehicle. n=4 independent experiments. Representative images of GLP-1 immunostaining (red) in organoids treated with (C) vehicle control and (D) 100μM indole. Nuclei are labelled by DAPI (blue). (E) L-cell numbers in control organoids and organoids treated with indole. n=27-30 organoids. (F) Blood glucose excursion during a IPGTT in C57BL6/J mice at baseline prior to treatment intervention.. n=8-10/group. (G) Blood glucose excursion during a IPGTT in C57BL6/J mice orally gavaged with 20mg/kg indole or water vehicle 15 minutes prior to glucose load (2g/kg). n=8-10/group.. n=8-10/group. (H) Blood glucose excursion during a IPGTT in C57BL6/J mice 3 days after a single oral bolus of indole. n=8-10/group. (I) Blood glucose excursion during a IPGTT in Pdx1-CreERT;Glp1r fl/fl mice 3 days after a single oral indole administration. n=6-7/group. Mice were orally gavaged with 20mg/kg indole or water vehicle and culled three days later. Representative images of ileal sections from (J) vehicle and (K) indole-treated mice following GLP-1 immunostaining (red). Nuclei are labelled by DAPI (blue). (L) The density of GLP-1 immunoreactive L-cells in the ileum of vehicle- and indole-treated mice.n=5/group. Insets in (F), (G), (H), (I) represent iAUC relative to t0. Scale bars in (C), (D), (J), (K) correspond to 50μm. (B), (E), (L) analysed by student’s t-test. (F)-(I) GTTs analysed by two-way ANOVA with Tukey’s post-hoc, iAUCs analysed by student’s t-test. Error bars represent SEM. *p<0.05, ***p<0.001, ****p<0.0001.

**Figure 6.**
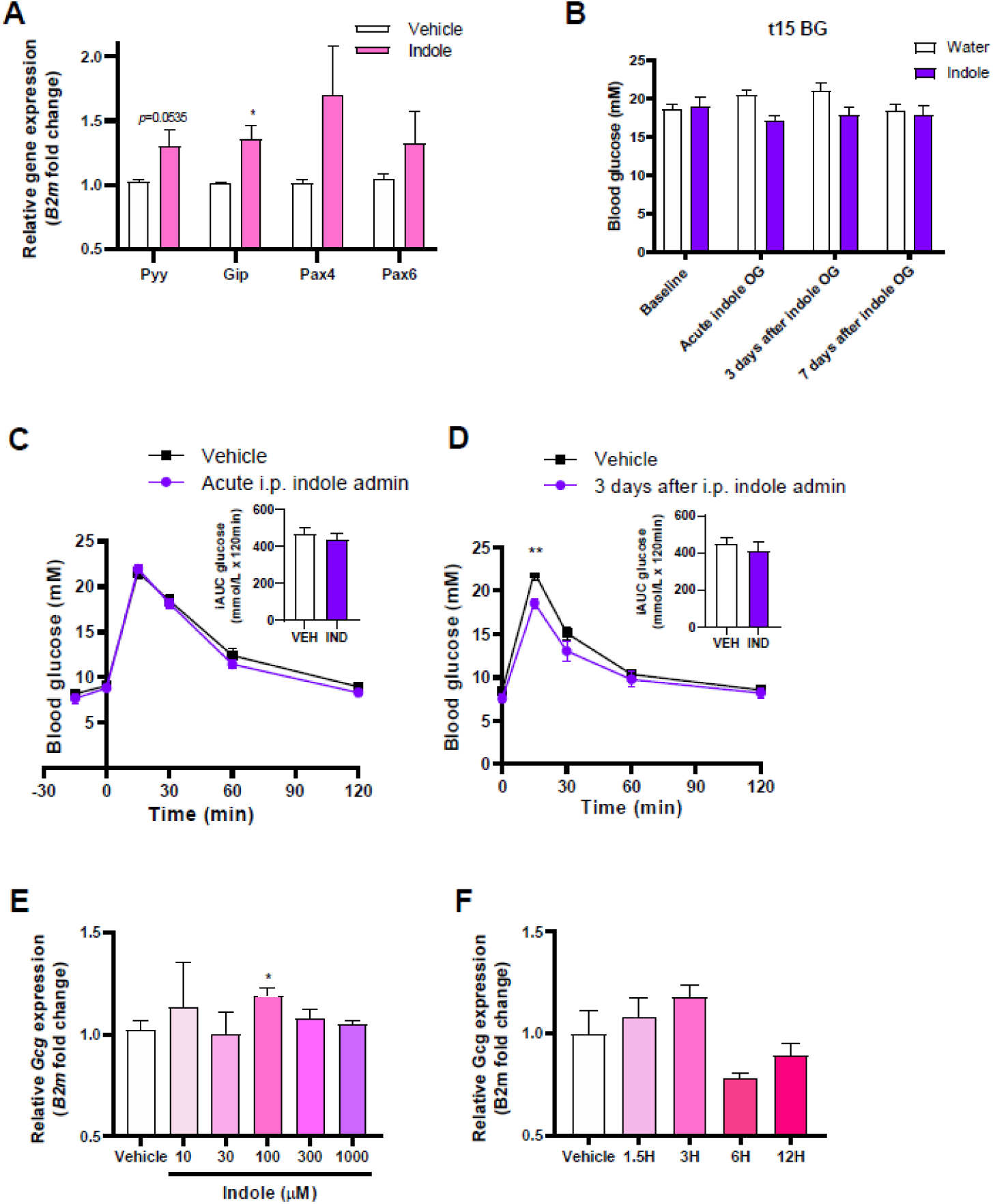
Further investigation of the effects of indole on enteroendocrine differentiation in intestinal organoids and in vivo. (A) Gene expression of the gut hormones Pyy and Gip, and the transcription factors directing L-cell development, Pax4, Pax6, in murine intestinal organoids cultured with indole or vehicle. n=4 independent experiments. (B) Blood glucose levels 15 min post intraperitoneal glucose (2g/kg) measured prior to indole administration, on the day of indole administration, 3, and 7 days after indole administration. n=8-10/group (C) Blood glucose excursion during a IPGTT in C57BL6/J mice that received an intraperitoneal injection of 20mg/kg indole or water vehicle 15 minutes prior to glucose load (2g/kg). n=8-10/group Inset represents iAUC determined relative to t0. (D) Blood glucose excursion during a IPGTT in C57BL6/J mice 3 days after a single intraperitoneal bolus of indole. Inset represents iAUC determined relative to t0. (E) The effect of 3h indole (10-1000μM) treatment on gene expression of the gut hormone Gcg in murine intestinal organoids. (F) The effect of 100μM indole treatment for 1.5-12h on gene expression of the gut hormone Gcg in murine intestinal organoids.

We next investigated whether the effects of indole on epithelial cell fate translated into metabolic benefits in vivo. Prior to a single oral bolus of indole treatment intervention, both groups of C57Bl6/J mice were similarly glucose tolerant (Fig 5F). Following acute oral indole administration, as expected, mice showed an improved glucose tolerance (Fig 5G). This improved glucose tolerance was sustained, with a reduced peak in blood glucose levels following an intraperitoneal bolus of glucose three days after oral indole administration (Fig 5H). This is despite the half-life of indole previously being reported to be less than 8 hours^21^. This effect of indole on glucose tolerance was lost by seven days post-administration (Fig 6B). Next, we observed that whilst intraperitoneal indole administration (which does not affect GLP-1 secretion) did not modify acute gluco-regulation (Fig 6C), 3 days post intraperitoneal administration, indole-treated mice displayed an improved glucose tolerance (Fig 6D). These data suggest the effect of indole on long-term glucose tolerance is mediated via a mechanism distinct from that mediating the acute stimulation of GLP-1 release. However, Pdx1-CreERT;Glp1r fl/fl mice lost these effects on glucose tolerance three days after oral indole administration (Fig 5I), suggesting that the mechanism for these longer effects improving glucose tolerance is also dependent on GLP-1 receptor signalling. These findings suggest that indole improves acute and long-term glucose tolerance via two distinct mechanisms, but that both involve GLP-1 signalling in the pancreatic beta cell.

The intestinal epithelium undergoes rapid renewal, regenerating every 3-5 days. Considering that the improvement in glucose tolerance observed in indole-treated mice was lost after approximately a week, and in line with the in vitro findings, we hypothesised that a single dose of indole would increase differentiation of GLP-1 secreting L-cells to improve glycaemic control. Ileum samples from C57Bl6/J mice that received a single oral indole dose collected three days later showed a ∼70% increase in L-cell density (Fig 5J-L).

Taken together, these data suggest that indole modulates EEC fate specification and increases L-cell abundance, improving glucose homeostasis in mice. These findings position indole as a potential anti-diabetic therapy.

### 3.3 Sub-chronic indole administration improves glucose tolerance and insulin sensitivity in WT and diabetic mice

To further investigate the therapeutic utility of indole in T2D treatment, we next studied the effects of sub-chronic indole administration in WT and T2D mouse models (Fig 7A). WT C57Bl6/J mice that received sub-chronic indole administration showed reduced fasting blood glucose levels (Fig 7C), in addition to improvements in intraperitoneal and oral glucose tolerance (Fig 7D, E); a modest increase in insulin sensitivity was also noted (Fig 7F). Notably, these effects were independent of any changes in body weight (Fig 7B). These data are in line with findings that chronic indole supplementation in mice reduced HFD-associated systemic insulin resistance and glucose intolerance, although this is the first study to report metabolic improvements after a relatively short treatment intervention, and in WT mice. Given these metabolic benefits of indole, and the preservation of GLP-1’s insulinotropic effect in T2D, the same sub-chronic indole treatment paradigm was repeated in TALLYHO/JnG mice, a polygenic T2D mouse model that exhibits hyperglycaemia and insulin resistance, reflecting the multifactorial nature of human T2D^30^. Indole-treated TALLYHO/JnG mice showed improved glucose tolerance (Fig 7G), and interestingly, a notable improvement in insulin sensitivity (Fig 7H), demonstrating the potential translatability of indole’s metabolic effects to a disease state. To further investigate this insulin sensitivity phenotype, we profiled whole liver transcriptomes from indole- and vehicle-treated TALLYHO mice. A total of 719 differentially expressed genes (unadjusted p<0.05) were found (Fig 7I), including an upregulation of Hk3, the rate-limiting enzyme of many glucose metabolism pathways, which represents a possible mechanism by which indole improves hepatic insulin sensitivity. Ingenuity pathway analysis further revealed that indole induces mild transcriptional alterations in liver metabolism; specifically, an inhibition of IL-6 cytokine signalling (Fig 7J-L). The anti-inflammatory effects of indole in the liver could potentially enhance insulin sensitivity too, consistent with work showing that indole and related compounds exert hepatoprotective effects in non-alcoholic fatty liver disease by reducing production of pro-inflammatory mediators^21,31,32^. Furthermore, modulation of cell-cell and tight junction assembly pathways were observed, in line with previous reports of indole’s role in maintaining epithelial barrier function (Fig 7K)^6,7^.

**Figure 7.**
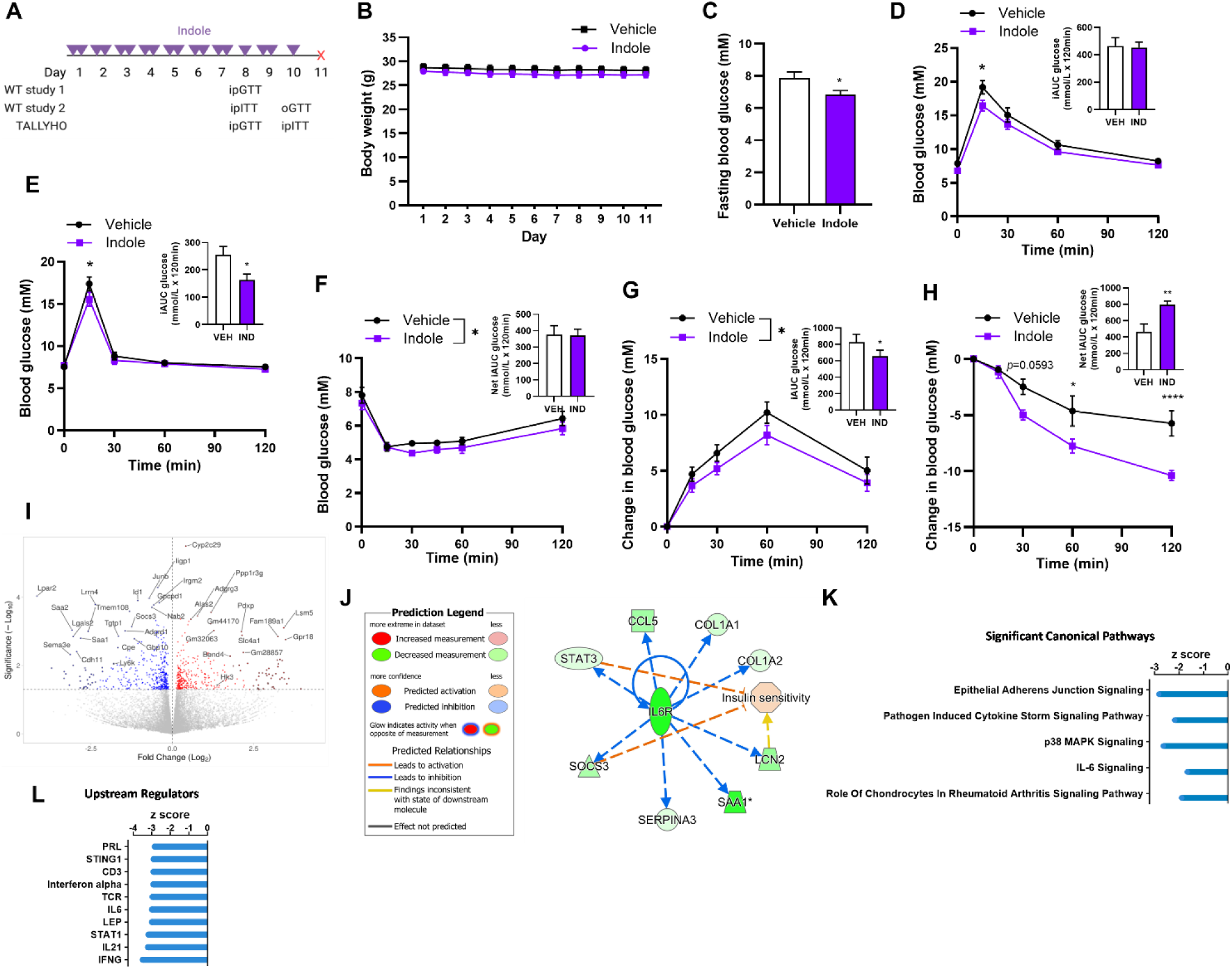
Sub-chronic indole administration improves glucose tolerance and insulin sensitivity in WT and T2DM diabetic mice potentially via inhibition of IL-6 signalling. (A) Sub-chronic indole administration study design. Mice received an oral gavage of 20mg/kg indole or vehicle control (water) twice daily for seven consecutive days. On day 8, an intraperitoneal insulin tolerance test (IPITT) or IPGTT was performed. The next day, mice received two doses of indole or vehicle. On day 10, an OGTT or IPITT was performed. One day after the second metabolic test, the mice were culled for tissue collection. (B) Body weight of vehicle- and indole-treated WT C57BL6/J mice over the course of the study. n=9-10/group. (C) Fasting blood glucose levels in WT C57BL6/J mice after seven days of indole or vehicle administration. n=9-10/group. Student’s t-test. (D) Blood glucose excursion during a IPGTT in vehicle- and indole-treated WT C57BL6/J mice. n=9-10 animals per group. (E) Blood glucose excursion during an OGTT (3g/kg glucose) after a 5h fast in vehicle- and indole-treated WT C57BL6/J mice. n=8-9/group. (F) Blood glucose excursion during an intraperitoneal insulin tolerance test (1IU/kg insulin) after a 5h fast in vehicle- and indole-treated WT C57BL6/J mice. n=8-9/group. (G) Change in blood glucose excursion (normalised to t0) during a IPGTT after an overnight fast in vehicle- and indole-treated diabetic TALLYHO mice. n=7/group. (H) Change in blood glucose excursion (normalised to t0) during a IPITT in vehicle- and indole-treated diabetic TALLYHO mice. n=7/group. (I) Volcano plot of differentially expressed genes between vehicle and indole-treated TALLYHO liver tissues. n=7/group. (J) Regulatory network for the IL-6 receptor overlayed with predicted effect on the IPA functional gene set for insulin sensitivity. Vertical Oval – transmembrane receptor, Horizontal oval – Transcriptional regulator, Triangle –phosphatase, Trapezoid – transporter, Square – Cytokine, Octagon – function, circle-other. (K) Z-score of significant (p<0.05) affected canonical pathways derived via Ingenuity Pathway Analysis (IPA) gene ontology analysis. (L) Top ten inhibited upstream regulators. Insets in (D), (E), (F), (G), (H) represent iAUC relative to t0. (D)-(H) GTTs and ITTs analysed by two-way ANOVA with Tukey’s post-hoc. Data in (C), and iAUCs in (D)-(H) analysed by student’s t-test. Error bars represent SEM. *p<0.05, **p<0.01, ****p<0.0001.

## 4. Discussion

We have shown that the bacterial tryptophan metabolite indole can act on the gastrointestinal tract to modify GLP-1 secretion from enteroendocrine L cells using several models. The acute stimulatory effect appears to involve the Trpa1 ion channel, but not the AhR, potentially by modulating extracellular Ca^2+^ flux. Importantly, we also demonstrate that the enhanced GLP-1 and hence insulin secretion translates into improved acute glucoregulation in vivo. Interestingly, indole potentially seemed to acutely worsen glucose tolerance in vivo when GLP-1R signalling was blocked, suggesting the beneficial actions of indole via GLP-1 may mask or prevent other less powerful deleterious effects.

A previous study reported time-dependent effects of indole on GLP-1 release from primary L-cells, with release suppressed at later time points^8^. We noted that longer treatment lengths or higher indole concentrations also reduced Gcg mRNA expression, suggesting that pulses of indole may more effectively improve glucose control than continuous exposure (Fig 6E,F). This is the first organoid study to investigate the effect of indole in modifying EEC specification. Interestingly, indole can increase neurogenesis in the adult mouse hippocampus^29^, and the transcriptional pathways implicated in gut endocrine specification and neuronal differentiation share some commonalities^33,34^. Nonetheless, future studies are required to elucidate indole’s effects on various intestinal cell types, as well as its transcriptional targets and regulatory pathways.

It is interesting to note that indole improved insulin sensitivity in healthy and diabetic mice, but that the effect was greater in the TALLYHO/JnG model of type 2 diabetes. This greater effect may reflect the anti-inflammatory effects of indole^21,31,32^, and suggests that indole may be useful in metabolic diseases associated with greater inflammatory signalling.

In conclusion, our work has identified indole as a novel anti-diabetic signalling molecule, providing further insight into its role in glucose control and L-cell differentiation. Further work is thus required to determine whether this pathway is conserved in humans, but our findings raise the novel possibility of incorporating indole into nutraceutical supplements treat and/or prevent glucose intolerance associated with T2D.

## Acknowledgments

The authors would like to thank Stephen Rothery of the Facility for Imaging by Light Microscopy at Imperial College London for his support and assistance in this work.

## Author contributions

PP: Conceptualisation, Investigation, Formal analysis, Project administration, Writing – original draft. MN: Methodology, Investigation. SC, AGR, LM, P-EC, CD, EO, SA: Investigation. RK: Investigation, Formal analysis. BYHL: Methodology, Visualisation, Formal analysis. FMG, FR: Resources, Writing – review & editing. ACH: Methodology, Writing – review & editing. GSHY: Methodology. GAB: Visualisation, Formal analysis. BJ: Methodology, Resources, Writing – review & editing. BO: Writing – review & editing, Methodology. KGM: Writing – review & editing, Supervision, Formal analysis, Data curation, Conceptualisation, Funding acquisition.

## Funding Sources

The Section of Endocrinology and Investigative Medicine is funded by grants from the MRC, BBSRC, NIHR and is supported by the NIHR Biomedical Research Centre Funding Scheme. The views expressed are those of the authors and not necessarily those of the MRC, BBSRC, the NHS, the NIHR or the Department of Health and Social Care. PP is funded by a National Science Scholarship from A*STAR, Singapore. KGM is supported by Diabetes UK (18/0005886, 20/0006295), the BBSRC (BB/W001497/1, BB/X017273/1), the MRC (MR/Y013980/1) and the Wellcome Trust (310835/Z/24/Z). BJ acknowledges funding support from the Medical Research Council (MR/Y00132X/1 and MR/X021467/1), the Wellcome Trust (301619/Z/23/Z), Diabetes UK, the Eli Lilly LRAP programme, and Metsera Inc. BYHL and GSHY are supported by BBSRC Project Grant (BB/S017593/1) and the MRC Metabolic Diseases Unit (MC_UU_00014/1). Next-generation sequencing was performed at the IMS Genomics and Bioinformatics Core supported by the MRC (MC_UU_00014/5, RGAG/542 MC_UU_00039) and the Wellcome (208363/Z/17/Z, RGAG/546 226800/Z/22/Z), and the Cancer Research UK Cambridge Institute Genomics Core. The FR/FG laboratories were funded by the Wellcome Trust (220271/Z/20/Z) and MRC (MRC_MC_UU_12012/3). The study’s funders were not involved in the design of the study; the collection, analysis, and interpretation of data; writing the report; and did not impose any restrictions regarding the publication of the report

## Disclosures

BYHL provides renumerated consultancy for Nuntius Therapeutics. GSHY receives grant funding from Novo Nordisk and Amgen Inc; he also consults for both Novo Nordisk and Eli Lilly and Company. The other authors declare no competing interests.

## Data Availability

All data, methods and unique/stable reagents generated in this study will be made available from the lead contact upon reasonable request, and may require a completed Materials Transfer Agreement.

